# Why Montane *Anolis* Lizards are Moving Downhill While Puerto Rico Warms

**DOI:** 10.1101/751941

**Authors:** C. J. Battey, Luisa M. Otero, George C. Gorman, Paul E. Hertz, Bradford C. Lister, Andrés García, Patricia A. Burrowes, Raymond B. Huey

## Abstract

Because Puerto Rico has warmed in recent decades, ectotherms there should have shifted their elevational ranges uphill. However, by comparing historical versus recent distributional records of *Anolis* lizards, we found that three “montane-forest” species have instead moved downhill in recent decades, almost to sea level. This downward shift appears related to the massive regeneration of Puerto Rican forests – especially in lowland areas – which started in the mid-20th century when the island’s economy began shifting from agriculture to manufacturing. The magnitude of local cooling caused by regenerated forests swamps recent climate warming, seemingly enabling cool-adapted “montane” lizards to track forests as they spread downhill from mountain refugia into abandoned plantations. Thus, contemporary distributional patterns are likely converging to those prior to the arrival of European settlers, who cleared most lowland forests for agriculture, thereby restricting forests and associated fauna to high-elevation remnants. In contrast with the montane species, three lowland species expanded their ranges to higher elevations in recent decades; but whether this movement reflects warming, land-use shifts, or hurricane-induced destruction of upland forests is unclear.

## 3 Introduction

A biogeographic shift to higher elevations is a common response of species to warming climates, at least where mountains are accessible (Colwell et al., 2008; Freeman and Freeman, 2014; Moritz et al., 2008; Parmesan, 2006). Such upward shifts enable organisms to track their thermal niche, but sometimes expose them to novel environmental conditions (Camacho et al., 2018) and species interactions (Angert et al., 2013). Over time the cumulative effect of these upward shifts may cause “biotic attrition” in lowland regions of the tropics, because the loss of lowland diversity caused by organisms shifting uphill is unlikely to be mitigated by new species moving in from lower latitudes (Colwell et al., 2008).

An upward range shift is, however, far from a universal outcome under warming (Lenoir et al., 2010; Lenoir and Svenning, 2015). In fact, numerous species have shifted ranges downhill, possibly reflecting competitive release (Davis et al., 1998), habitat or land use changes (Guo et al., 2018; Santos et al., 2017), or influences of non-thermal aspects of climate (Angert et al., 2013; Lenoir et al., 2010; McCain and Colwell, 2011).

Most evaluations of elevational range shifts have focused on temperate species (Chen et al., 2011; Guo et al., 2018; Lenoir et al., 2010), mainly because these species usually have more complete distributional records. Even so, tropical ectotherms might show conspicuous elevational responses to warming because (1) their ecology is tightly coupled to environmental temperature, (2) their physiology is often specialized for temperature (Deutsch et al., 2008; Polato et al., 2018), (3) their elevational ranges are narrow (Huey, 1978; Wake and Lynch, 1976; Polato et al., 2018), and (4) tropical thermal regimes shift rapidly with elevation (Janzen 1967, but see Buckley 2013). Nevertheless, comparatively few studies have evaluated range shifts on tropical mountains (but see Freeman and Freeman 2014; Guo et al. 2018; Lenoir et al. 2010; Morueta-Holme et al. 2015; Pounds et al. 1999; Raxworthy et al. 2008).

*Anolis* lizards on the mountainous tropical island of Puerto Rico provide an opportunity to explore elevational range shifts over the last half century, because their distributions, behaviors, ecologies, and thermal biologies are well known (Gorman and Licht, 1974; Gunderson and Leal, 2012; Hertz et al., 1979; Huey and Webster, 1976; Leal and Fleishman, 2002; Lister, 1981; Losos, 2011; Rand, 1964; Rodríguez-Robles et al., 2005; Schoener, 1971; Williams, 1972). Moreover, Puerto Rico has warmed slightly in recent decades (around 0.3°C in our analysis, but as much as 2.4°C in other analyses (Burrowes et al., 2004; Comarazamy and González, 2011; Jennings et al., 2014; Méndez-Lázaro et al., 2015; Méndez-Tejeda, 2017; Waide et al., 2013; Lister and García, 2018), thus setting an expectation for ranges of montane species to have shifted uphill.

We have been studying responses of Puerto Rican anoles to recent climate warming (Lister and García, 2018), and one of our initial planned projects was to evaluate whether the lower elevational range limit of *Anolis gundlachi* had shifted uphill in recent decades. This lizard has long been described as a montane, forest-dwelling thermoconformer (Gorman and Licht, 1974; Heatwole, 1970; Hertz et al., 1979; Hertz, 1981; Huey and Webster, 1976; Lister, 1981; Rand, 1964; Rivero, 1998; Schmidt, 1918; Schoener, 1971; Williams, 1972). Relative to other Puerto Rican anoles, it is active at low body temperatures, is intolerant of high body temperatures, has high rates of evaporative water loss (Gorman and Hillman, 1977; Gunderson and Leal, 2012; Heatwole, 1970; Hertz et al., 1979; Hertz, 1981; Huey and Webster, 1976; Rand, 1964), and avoids sunny habitats and perches (Hertz, 1992; Rodríguez-Robles et al., 2005; Schoener, 1971). Therefore, shifts in its lower range limit should be a sensitive indicator of climate warming. Moreover, warming temperatures should enable *A. cristatellus* (a warm-adapted congener that inhabits forests in the lowlands as well as open habitats there) to invade mid-elevation forests from adjacent open habitats, adding competitive pressure on resident *A. gundlachi* (Buckley, 2013; Huey et al., 2009).

Extensive field research through the 1980s placed the lower limit of *A. gundlachi*’s elevational range as *≈* 200 to 250 m (Huey and Webster, 1976; Rivero, 1998; Schwartz and Henderson, 1991; Williams, 1972). Given an estimated temperature lapse rate of *-*6.5 ° C/km and an average increase in temperatures of 0.31°C (see Results), a naïve tracking model predicts the range of *A. gundlachi*would have shifted upward by approximately 46 meters. In 2011, we started a pilot study to determine if this prediction would hold. We drove down P.R. Highway 191 through Luquillo National Forest, stopped every 50-m drop in elevation, and searched adjacent forest for *A. gundlachi*. We kept finding this species at elevations below 250 m, and even found it as low as 20 m, adjacent to the floodplain of the Río Grande Luquillo! Thus, *A. gundlachi* appeared to have moved downhill since the 1970s, completely contrary to our expectations.

Our 2011 observations forced several post hoc questions concerning historical shifts in the distributional patterns of Puerto Rican anoles. First, were our brief surveys on Highway 191 correct and general for *A. gundlachi*? If so, records of *A. gundlachi* at other lowland localities would appear recent; and this could be evaluated by comparing historical versus contemporary museum and locality records. Second, do recent range shifts of other species of montane anoles match those of *A. gundlachi*? If so, elevation ranges of these species should also show similar descending trends. Finally, if both patterns hold, why would elevation ranges have expanded downward when temperatures were rising?”

Our efforts to answer the last question led us to the literature on historical changes in forest cover in Puerto Rico. Although forest cover has been long declining in many parts of the tropics and elsewhere (Taubert et al., 2018), forest cover in Puerto Rico has increased markedly since the middle of the 20th century (Álvarez-Berríos et al., 2013; Helmer et al., 2008; Lugo and Helmer, 2004), reflecting a shift from an agrarian rural economy to a manufacturing urban one (Grau et al., 2003; Rivera-Collazo, 2015; Yackulic et al., 2011). Abandoned plantations became re-forested during this period (Helmer et al., 2008) despite massive forest blowdowns caused by Hurricanes Hugo (1989) and Georges (1998).

Here we analyze historical versus recent patterns of locality records (largely museum collections) for the six most common species of Puerto Rican anoles. These records and other data presented here suggest that three montane species have been moving downhill since the mid-twentieth century, while three lowland species have expanded slightly uphill. We propose that the most plausible explanation for our findings is that regenerated forests have cooled and humidified lowland environments sufficiently to enable montane anoles, which are sensitivity to high temperature and desiccation, to re-invade the lowlands (see Discussion), which they likely occupied before the 16th Century when Europeans began clearing lowland forests (Lugo et al., 1981).

## 4 Methods

### 4.1 Study Species

The six common *Anolis* species studied here (*A. cristatellus*, *A. evermanni*, *A. gundlachi*, *A. krugi*, *A. pulchellus*, and *A. stratulus*) are members of a clade that evolved in situ on the greater Puerto Rican bank (Helmus et al., 2014). They are often partitioned into three pairs of “ecomorphs,” each of which has characteristic associations between microhabitats occupied and morphology (Williams, 1972). One member of each pair typically has a more lowland distribution. Even so, each pair of ecomorphs is broadly sympatric. Where sympatric, the lowland species is found in warmer, more open habitats and has higher body temperatures (Hertz et al., 2013; Huey and Webster, 1976; Rand, 1964), higher heat tolerance (Gorman and Licht, 1974; Gunderson et al., 2016; Huey and Webster, 1976), lower cold tolerance (Heatwole et al., 1969), and greater desiccation resistance (Gorman and Hillman, 1977; Hertz et al., 1979).

The two “trunk-ground” ecomorphs (*A. cristatellus* and *A. gundlachi*) typically perch low on shrubs and trees, foraging there or on adjacent ground. *Anolis cristatellus* has higher body temperatures, higher heat tolerances, and lower rates of evaporative water loss: it is found from sea level to high elevation. In the lowlands it occupies both forest and open habitats, but at mid- to high-elevations it is found only in open habitats (Huey, 1974; Huey and Webster, 1976; Otero et al., 2015; Schoener and Schoener, 1971). *Anolis gundlachi* has long been considered a “montane forest specialist” and restricted to deeply shaded, upland forests because of its sensitivity to high temperatures and its elevated rates of evaporative water loss (Gorman and Licht, 1974; Heatwole, 1970; Hertz et al., 1979; Huey and Webster, 1976; Lister, 1981; Rand, 1964; Rivero, 1998; Schmidt, 1918; Aide et al., 1996; Schoener, 1971; Williams, 1972).

Two “grass-bush” anoles (*A. pulchellus* and *A. krugi*) occur over broad elevational ranges. *Anolis pulchellus* is abundant at low to moderate elevation in relatively exposed habitats. *Anolis krugi* is more of a montane species but was sometimes found at low elevation “under conditions of extreme shade” (Gorman and Licht, 1974). The two “trunk-crown” anoles (*A. stratulus* and *A. evermanni*) perch somewhat higher in trees than the other species (Schoener, 1971). *Anolis stratulus* is broadly distributed, whereas *A. evermanni* is more of an upland form, found at sea level only in densely shaded habitats (Gorman and Licht, 1974).

### 4.2 Specimen Records and Georeferencing

We downloaded all specimen records of Puerto Rican *Anolis* from the Global Biodiversity Information Facility (GBIF) and developed scripts to remove duplicate records and apply consistent formatting across museums. Some records were geo-referenced, but many were not. Consequently, we manually georeferenced all non-GPS collection localities that had at least 10 specimens. We used Google Maps to measure road kilometers (many sites were listed as 10 km SE of a given town) and Google Earth to locate point localities. We followed MANIS best-practices for georeferencing (Wieczorek, 2001) and estimated uncertainty for all points to the nearest 100 m.

Because many surveyed areas were visited by multiple surveyors, often reporting slightly different GPS coordinates or text locality descriptions, we used a hierarchical clustering approach to group adjacent collecting sites into “locality clusters” which could be analyzed as independent geographic observations when testing for shifts in diversity. We calculated pairwise geographic distances among all unique specimen localities, applied a complete-linkage hierarchical clustering algorithm, and extracted clusters of localities within approximately 1 km of each other. We also added sight records made by us or by colleagues experienced with Puerto Rican *Anolis*.

After removing localities with greater than 2-km georeferencing uncertainty, we estimated the elevation of each collecting locality as the mean elevation across the full uncertainty radius in the USGS National Elevation Database (USGS, 2016) at 1-arc-second resolution, using the “rgeos” and “Raster” packages in R (Bivand and Rundel, 2013; Hijmans, 2015; Team, 2014). When surveyors reported an altitude directly, we used their data rather than a location-based estimate. Specimen records were binned into four time periods (see below) bracketing 1935-2015, matching the forest-age ranges used in Helmer et al.’s (2008) analysis of land-cover change on Puerto Rico. We subset records to include only the six most common species of *Anolis* (above), all of which occur in the mountains. The final dataset includes 8,839 specimens and 121 sight records across 505 localities and 293 locality clusters (Figure 1). Most collecting occurred during the periods 1952–1977 or 1991–2016 (Figure 1), so we compare these two time periods.

**Figure 1.**
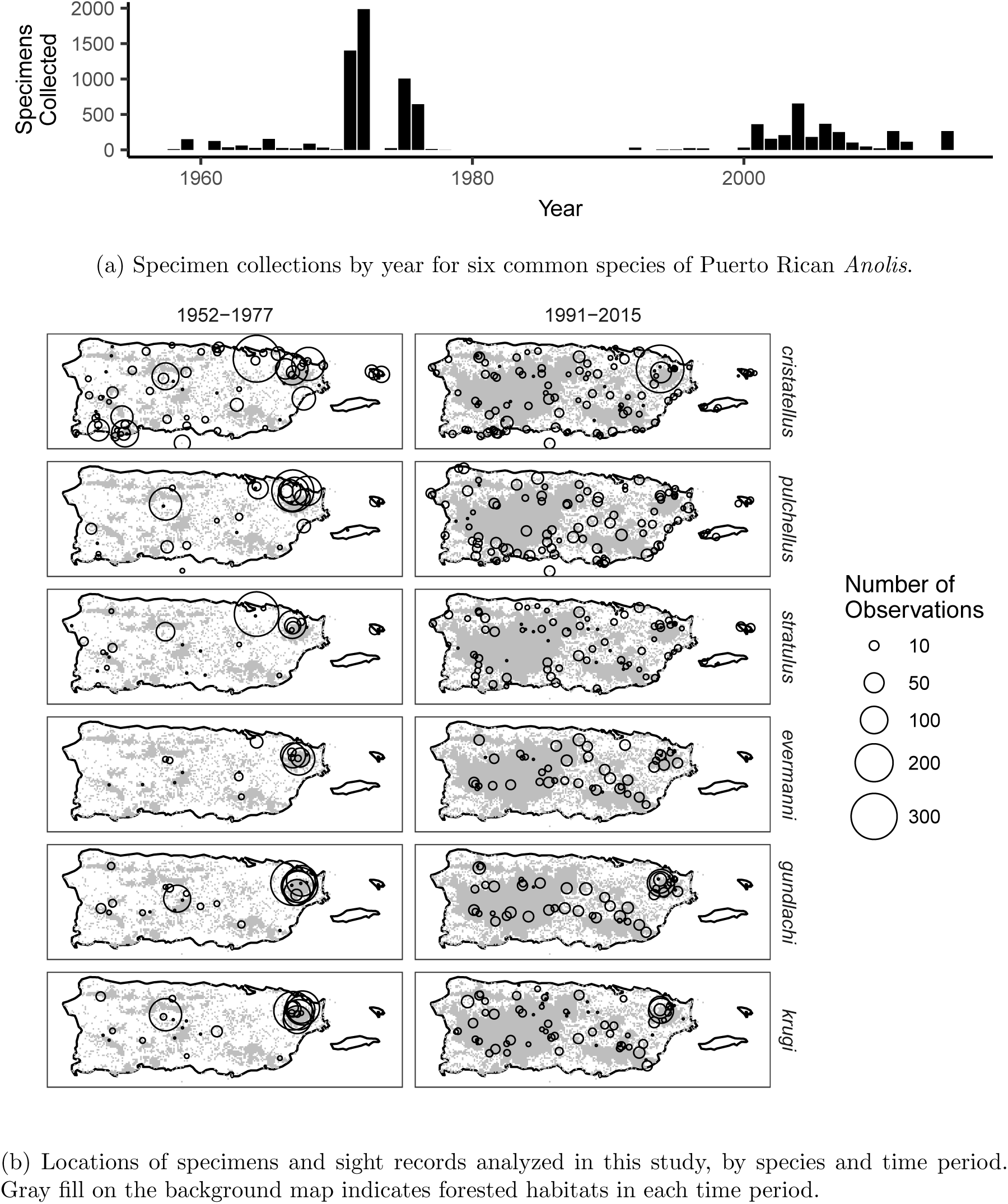

### 4.3 Shifts in Elevation Distribution

We used two complementary analyses to assess the extent and direction of elevational range shifts: a nonparametric hypothesis test that assesses the significance of shifts in median elevations across time periods, and a Bayesian linear model that estimates credible intervals while controlling for variation in survey effort by elevation. In the first analysis we determined the absolute elevation range of each species during each time period, then used a Wilcoxon rank sum test to evaluate whether the median elevation of specimen collections had shifted from 1952–1977 to 1991–2015. Because such shifts might merely reflect elevational shifts in collecting or survey effort between time periods, we set the null hypothesis for each species as the median difference in collecting elevation for all species other than the focal species. Thus, our test addresses the question “did the median elevation of species *i* shift more than expected, given overall shifts in *Anolis* collections?”

In the second analysis we modeled the abundance of species *i* across elevations as lognormally distributed, with mean equal to the sum of (1) the mean elevation for 1952-1977 occurrences (*A* _*i*_), (2) the difference caused by variation in survey effort across elevations (*E* _*i*_), and (3) the difference caused by true historical shifts in elevation distributions (*D* _*i*_). The log-normal was used because it forces positive values and mimics a decrease in total land area as altitude increases.

We set normally distributed priors for *A*, *E*, and *D*. *A* _*i*_ was parameterized with the empirical mean (*µ* _*i*_) and standard deviation (*sd* _*i*_) of 1952-1977 occurrences. For *E* _*i*_ we set the mean (*e* _*i*_) to the mean difference in collecting elevation across time periods for all species other than the focal species. The standard deviation (*s* _*i*_) was estimated by taking 100 bootstrap samples from the set of all non-focal-species reports and calculating the standard deviation of the differences in means. The mean of *D* _*i*_ was set to 0 and the standard deviation to 50 (we also ran the model with this value set to 200 and found minimal differences in results). The full model for species i is then:

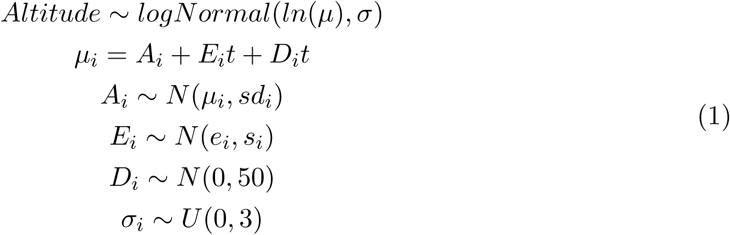

Where *t* is 0 for time period 1952-1977 and 1 for time period 1991-2015. We fit this model independently for each species using the maximum a posteriori (map) function from the R package “rethinking” (McElreath, 2012), and estimated credible intervals by sampling from the posterior. We note that an ideal analysis for the questions we addressed here would be an occupancy modeling approach similar to that described in Tingley and Beissinger (2009); however, our dataset lacks the repeated per-site sampling events (at least at most localities) required for this approach.

To assess potential changes in species composition at low elevations, we extracted specimen reports at elevations lower than 250 m, split them by species, and used a McNemar test to compare the relative abundance of each species across time periods (i.e., the number of specimens of species *i* relative to the number of all Anolis specimens in each period). This procedure is conceptually similar to the specimen-derived abundance index used in Linck et al. (2016) and Rohwer et al. (2012), in which the observed abundance of a target species is corrected for survey effort by dividing by the number of specimens collected with similar techniques in a given area. For all hypothesis tests conducted separately on each species, we corrected *p*-values for multiple testing (n=6) with the Holm-Bonferroni method (Holm, 1979).

Finally, to test if elevation range shifts are leading to loss of species diversity at low altitudes (see Colwell et al. (2008)), we calculated Shannon diversity (Shannon, 1948) for each locality cluster, dropped sites with only one reported species (because these may represent collections targeted at single species), and tested for differences in the distribution of species diversity of locality clusters across time periods in 250-meter elevation bins with a Wilcoxon rank-sum test. We chose the Shannon index as opposed to raw species richness because it is subject to less bias in the presence of undersampling (Beck and Schwanghart, 2010). To evaluate whether the arbitrary binning design (above) biased our results, we also fit linear models relating diversity to altitude (as a continuous variable) both with and without a categorical explanatory variable for time period, then used an ANOVA to test whether adding age class significantly improved model fit.

### 4.4 Changes in Land Cover and in Temperature

To assess changes in forest cover on Puerto Rico over the twentieth century, we modified an existing raster layer of forest age and soil types across the island developed from analysis of aerial and satellite imagery (Helmer et al., 2008). We merged forest ages across soil types and subset the original raster layer to produce maps of forested areas at 30-m resolution in four time bins: 1935–1951, 1952–1977, 1977–1990, and 1991–2000 (age classes in Helmer et al. (2008)). We then extracted the elevations of forested grid cells in each time period and calculated the total area and proportion of forested land in each 10-m elevation band across all time periods (Figure 3).

Several studies have documented increases in Puerto Rican temperatures during the late 20th Century (Burrowes et al., 2004; Comarazamy and González, 2011; Jennings et al., 2014; Méndez-Lázaro et al., 2015; Méndez-Tejeda, 2017; Waide et al., 2013; Lister and García, 2018), and average air temperatures on the island are predicted to continue to increase (Harmsen et al., 2009; Patz et al., 1998). To determine whether local warming is consistent with global observations and global climate model predictions (Karl et al., 2015), we examined temperature data (1950-2015) from the NOAA National Climate Data Center (https://www.ncdc.noaa.gov/cdo-web/search). Because change in precipitation could affect forest cover, we also examined change in total annual precipitation at these stations.

Eight weather stations on Puerto Rico reported at least 40 years of complete monthly data starting in 1950: Roosevelt Roads, Rio Piedras Experimental Station, Borinquén Airport, Lajas Substation, Manatí, Corozal, Ponce 4E, and Dos Bocas. Six of these stations are distributed around the periphery of the island near sea level, whereas two are in interior valleys at elevations between 100 and 200 meters (Appendix S1: Fig S4). For each of the eight stations, we removed any years without 12 months of data, calculated average annual temperatures, and used a t-test to compare the average temperatures between 1952–1977 and 1991–2015. To avoid biases in temperature records caused by the urban heat island effect (Oke, 1982; Murphy et al., 2011), we identified stations in urban areas as of 1991 by extracting the land cover class from Helmer et al.’s 2008 forest map. Finally, to estimate the rate of temperature change with altitude, we fit a linear model for average annual temperature across all years as a function of altitude, using a larger set of 16 weather stations (including those with fewer than 40 years of data) spanning 4 to 1144 m in elevation.

Full datasets and scripts for all analyses are available at: https://github.com/cjbattey/anolis elevation shift.

## 5.Results

### 5.1 Shifts in Elevation Ranges

Analyzed on a per-occurrence basis, median elevations for all of the six common *Anolis* species on Puerto Rico shifted significantly between 1952–1977 and 1991–2015 (Figure 2, Table 1). In general, highland species expanded downhill, while lowland species expanded uphill. *Anolis gundlachi* showed the largest decreases both in median elevation (480 to 385 m) and lower elevation limit (223 to 24 m). *Anolis cristatellus* had the largest increase in maximum elevation; rising from 880 m in the mid-20th century to 942 m after 1991.

**Table 1:**
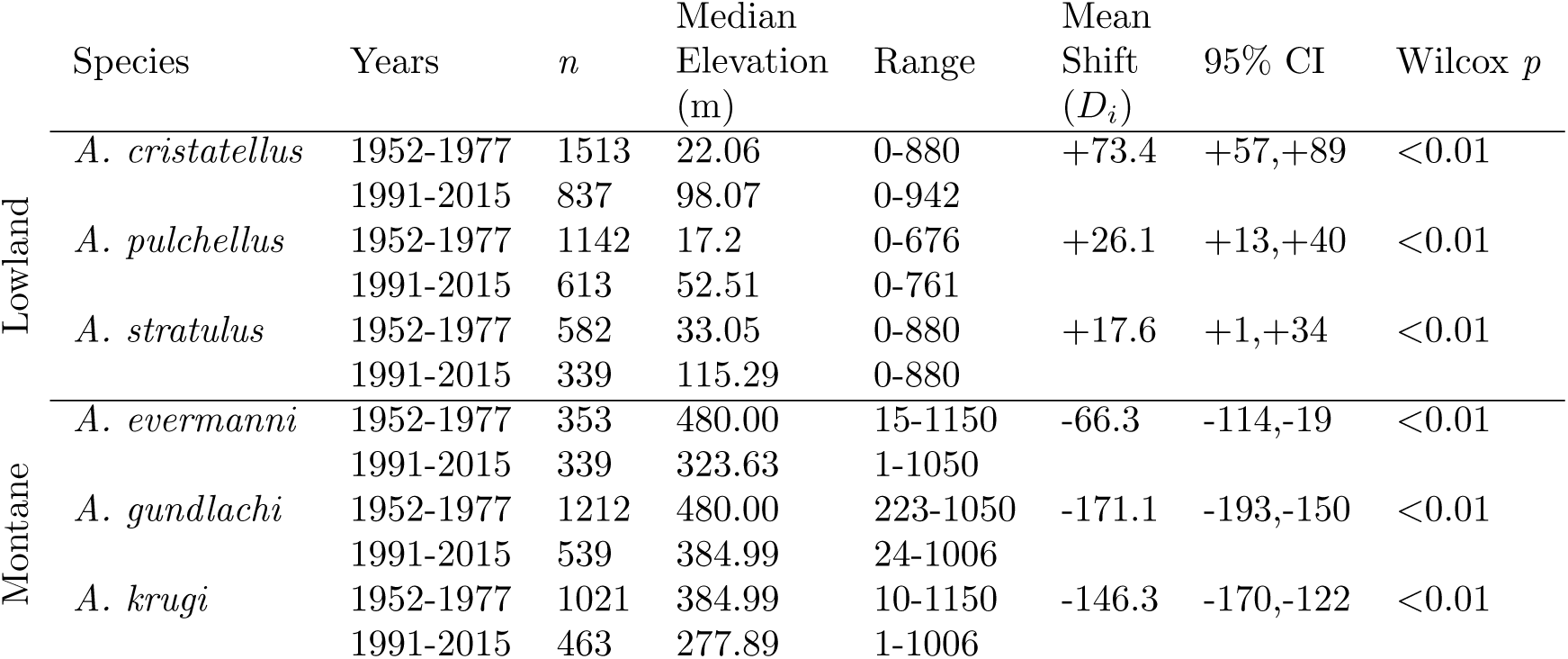
Summary of elevation ranges by species and time period. The mean and 95% CI describe the estimated elevation shift in meters while accounting for differences in survey effort across time periods. *P* values are for differences in medians in a Wilcox test.

**Figure 2:**
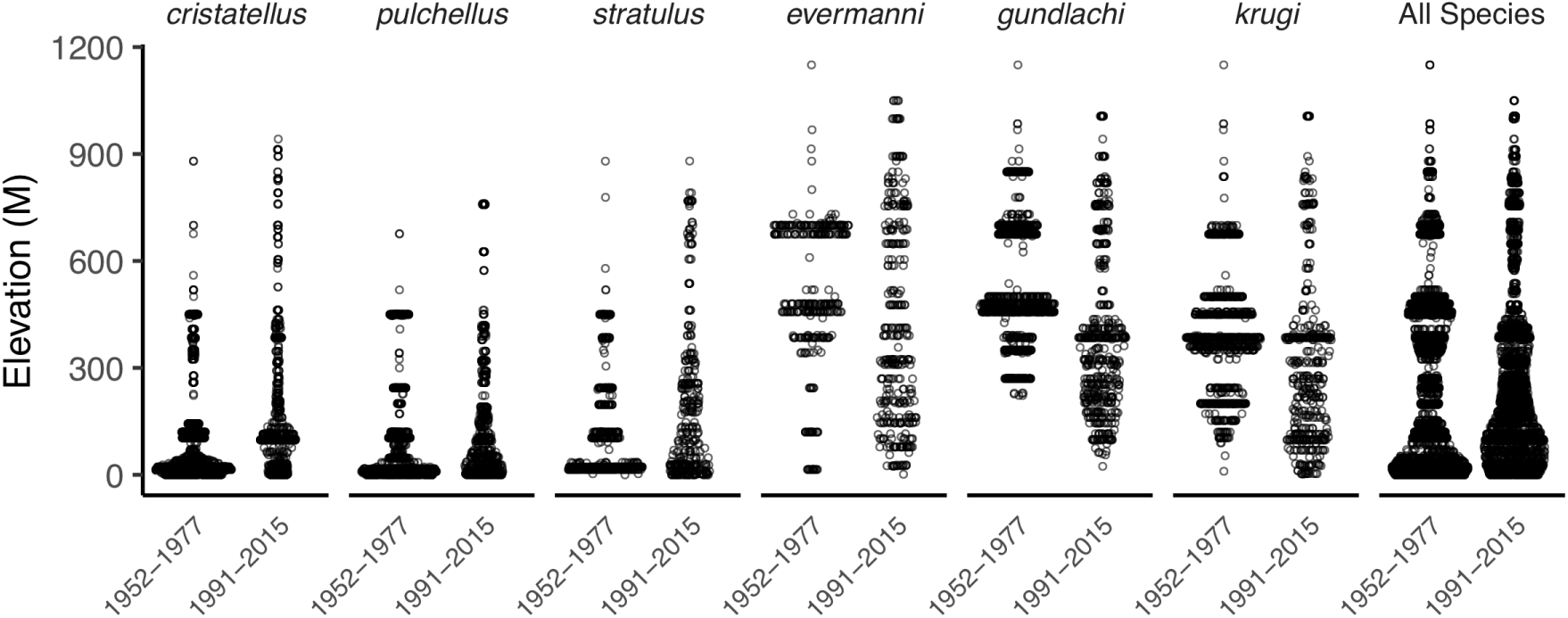
Elevation distributions for six common species of Puerto Rican *Anolis* in the periods 1952-1977 and 1991-2015. Points are spread horizontally proportional to the density of occurrences at a given elevation. The first three species (from left to right) are lowland, and the rest are montane.

Credible intervals in Bayesian models for the extent of the historical range shift suggest that differences in the elevational distribution of survey effort across time periods do not fully explain the observed shifts in any species (Table 1, Table S1, Figure S1). The largest downward shift in mean elevations was estimated in *A. gundlachi* (−171 meters), whereas the largest uphill shift was estimated in *A. cristatellus* (73 meters). This is consistent with our analysis of relative frequencies at sites below 250 m, where we observed that “lowland” species became less common and “montane” species more common (Table 2).

**Table 2:**
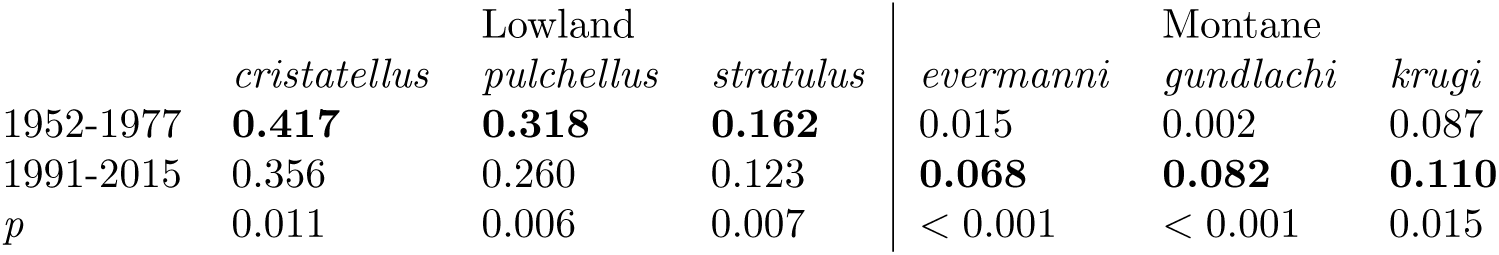
Frequency of each species as a proportion of all *Anolis* occurrences below 250 meters, with holm-corrected p values from a chi-squared test for change between periods. The higher frequency is bolded for each species.

Comparing Shannon diversity of locality clusters in elevation bins across time periods, we found no evidence of a loss of lowland diversity over time (Colwell et al., 2008). Observed lowland diversity (elevations below 250 m) had increased between time periods, but the median shift was not statistically significant (*p* = 0.066, Table 3, Figure S2). In the linear modeling framework, diversity is weakly but positively correlated with elevation (*intercept* = 0.72, *slope* = 0.0004, *p* (*slope >* 0) = 0.02, *R* ^2^ = 0.049), but adding time period as a categorical explanatory variable did not significantly improve model fit (anova *df* = 2, *p* = 0.226). Thus both analyses suggest that the relationship between diversity and altitude has not changed significantly between time periods, despite shifts of individual species.

**Table 3:** Difference in Shannon diversity of locality clusters by elevation and time period, with *p* values from a Wilcoxon rank-sum test.

| Elevation<br>Range (m) | Years | *n* locality<br>clusters | Median<br>Diversity | Shift | *p* |
| --- | --- | --- | --- | --- | --- |
| 0-250 | 1952-1977 | 23 | 0.683 | 0.115 | 0.066 |
|  | 1991-2015 | 83 | 0.783 |  |  |
| 250-500 | 1952-1977 | 13 | 1.040 | 0.126 | 0.423 |
|  | 1991-2016 | 25 | 1.100 |  |  |
| 501-750 | 1952-1977 | 6 | 0.818 | 0.025 | 0.743 |
|  | 1991-2015 | 12 | 1.015 |  |  |
| 751-1150 | 1952-1977 | 5 | 1.004 | 0.065 | 0.853 |
|  | 1991-2015 | 14 | 1.017 |  |  |

### 5.2 Land Cover and Temperature

Forests in Puerto Rico expanded dramatically by almost five-fold (as a percent of total land area) from 1935-1951 (8.9%) to 1991-2000 (43.1%) (Helmer et al., 2008). All elevation bands showed significant increases in forested area, but elevations of 100-300 M had the largest absolute increases (Figure 3B). In total roughly 1,200 square kilometers of land area converted from open to forested habitats in this single elevation range from 1935 to 2000. Forests expanded across the island, but the effect was particularly marked in the west (Figure 3A).

**Figure 3:**
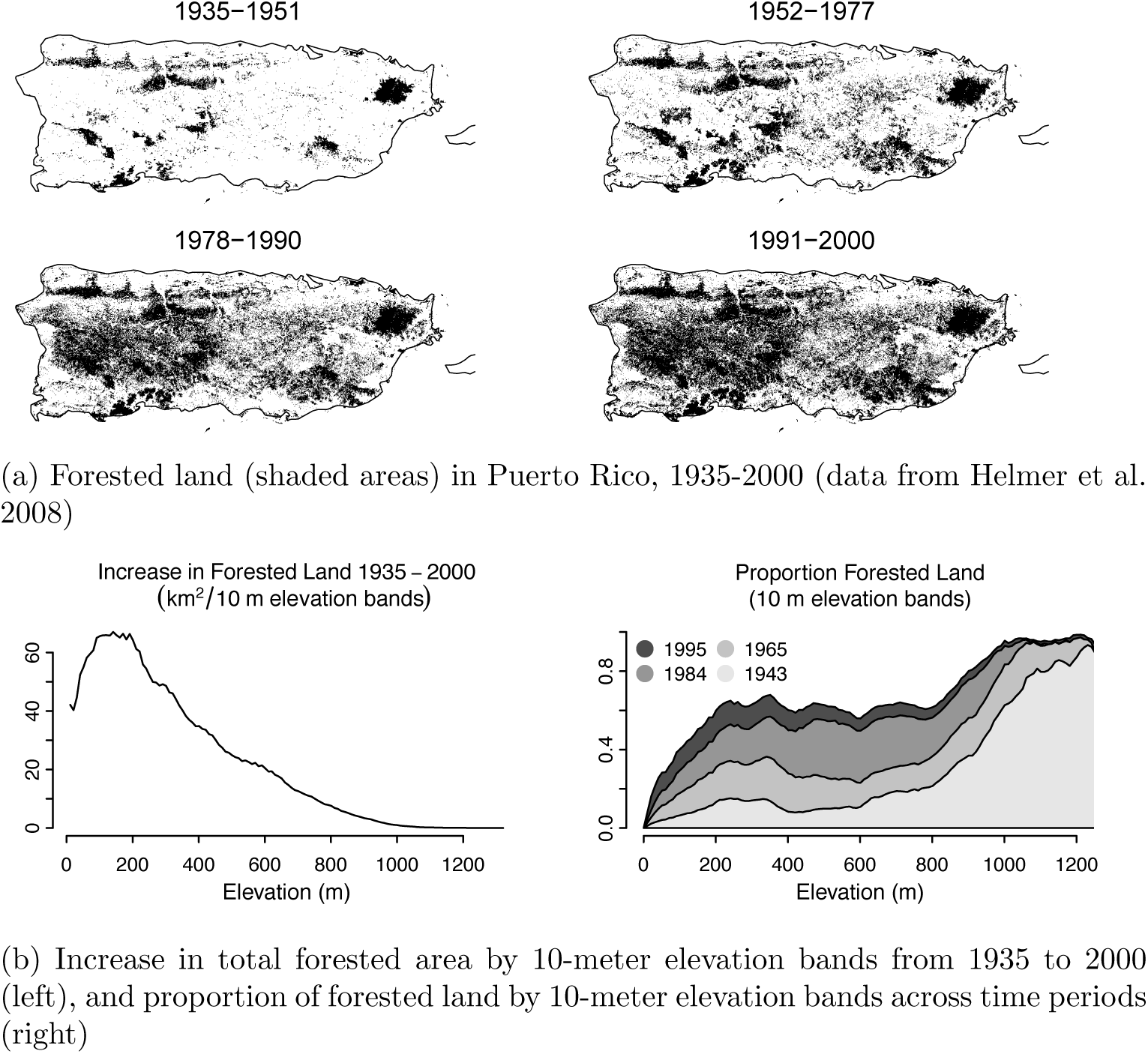
Forest regrowth in Puerto Rico.

Average annual temperatures increased significantly between 1952–1977 and 1991–2015 at four of eight weather stations (4; Figure S3), with an average shift across all sites of 0.31°C. One site (Dos Bocas, 119 m) was cooler after 1991, but all other sites warmed. If sites in urban areas are removed, the average change in temperatures drops to 0.22°C. Mean annual precipitation differed significantly at two of eight stations (Appendix 1: Table S2, Fig S4), with an average change across all sites of −14.46 cm per year (approximately a 10% decrease relative to the 1952-1977 average). The estimated temperature lapse rate across our expanded set of 16 short-term stations was −6.5°C/km (p*<* 0.001, *R* ^2^=0.887, Figure S4).

The only other study we are aware of comparing temperature records from ground stations on Puerto Rico over a similar timespan is Méndez-Tejeda (2017), which concluded that average temperatures had increased by 2.24°C between 1950 and 2009. Unfortunately, we were unable to determine the exact methods used to arrive at this figure and so cannot comprehensively explain the difference with our result. However, we note that the 2.24°C is also significantly higher than the 0.6-0.9°C increase in annual temperatures between 1950-1959 and 2000-2004 estimated for the area around Puerto Rico in the Caribbean-wide study by Comarazamy and González (2011).

Although no long-term data were available for high-elevation sites, one study (Burrowes et al. (2004), at about 1000 m) recorded an average minimum temperature increase of 0.72°C from 1970 to 2000, suggesting that temperature increases might have been higher in montane areas than in lowland area. Lister and García (2018) also estimated a roughly 1.5°C increase in monthly *maximum* temperatures at two sites near 350 M in elevations from c. 1980-2010. Thus, although significant uncertainty remains about the magnitude of temperature increases on Puerto Rico (particularly at higher elevations), all studies agree that temperatures have increased.

Because *A. gundlachi* is a thermoconformer, its body temperature is close to that of ambient air temperature in shaded areas (average absolute deviation 0.4°C, see Huey and Webster 1976). If its lower range limit is set by temperature, and if recent warming was 0.3°C, then the lower limit should have shifted up by approximately 46 m, given a lapse rate of −6.5°C/km (above)).

## 6 Discussion

Before we started our field work, we expected that elevational ranges of “montane” species of *Anolis* lizards (especially *A. gundlachi*) in Puerto Rico would have shifted upward in response to observed recent warming (Burrowes et al., 2004; Jennings et al., 2014; Méndez-Lázaro et al., 2015; Waide et al., 2013). To our surprise, our ground surveys – and the subsequent analysis of specimen records described above – showed just the opposite: all three of the montane species now occur at lower elevations than they did in the midtwentieth century (Figure 2), and *A. gundlachi* – the least heat tolerant of these species – is now found even near sea level.

Are these findings an artifact of collecting bias? To be sure, many of the collections were never intended as general biodiversity surveys and lack detailed information on survey effort and on species inventories. Although we controlled for differences in the elevation distribution of survey effort across time periods (see Methods), we cannot exclude the possibility that some surveyors concentrated preferentially on different species in different time periods or areas. For example, perhaps surveyors found but did not collect *A. gundlachi* in low-elevation forests in the mid-20th century, but started collecting this species there only recently. However, based on our personal histories of fieldwork on Puerto Rico (collectively the authors recorded 34% of all occurrences in the dataset, including 52% of those in the 1952-1977 time period), we feel confident that the absence of montane species in records at low elevations during the mid-20th Century is correct and that our core result – downward shifts of montane species – is not caused by systematic biases in surveyor behavior.

What then explains the elevation shifts we observed in montane *Anolis*? The most plausible explanation relates to historical changes in land use in Puerto Rico. In many tropical regions, logging and agriculture have reduced forest cover (Álvarez-Berríos et al., 2013; Guo et al., 2018). But in Puerto Rico forests have expanded substantially (Fig. 2) since the middle of the 20th Century (Helmer et al., 2008; Lugo and Helmer, 2004), reflecting a shift from an agricultural and rural economy to a manufacturing and urban one (Yackulic et al., 2011), and with abandoned farms converting to forests.

Such a rapid expansion (Figure 3) may appear surprising, but forest recovery following hurricanes is much faster in the tropics than in temperate zones (Canham et al., 2010). Indeed, Puerto Rican forests recover basic structure within a decade or two after hurricane blowdowns (Lugo and Helmer, 2004; Walker, 1991), and are indistinguishable from primary forest in terms of density and tree size after only 40 years of recovery (Aide et al., 1996).

Although deforestation can accentuate the biological impact of climate warming (Guo et al., 2018), the successional replacement of an open habitat by a forest acts antagonistically to climate warming and reduces (in the understory) incident radiation, maximum air temperature, and maximum wind speed, while raising relative humidity (Bastable et al., 1993; Geiger et al., 2009). Consequently, “operative” temperatures (a thermal index of the equilibrium body temperatures of a dry-skinned ectotherm, see Bakken (1992)) will be much cooler in the forest understory than in adjacent open habitats (Kaspari et al., 2015; Otero López and Sabat, 2015). For example, at a lowland site in Puerto Rico in summer (Monagas, see Otero et al., 2015), operative temperatures in the open often exceeded 40°C (maximum = 46.4°C), which is well above the critical thermal maximum of *A. gundlachi* (37.5°C, Huey and Webster (1976)), whereas operative temperatures in the adjacent forest rarely exceeded 30°C (maximum = 33.3°C).

The magnitude of understory cooling driven by a forest is thus much larger than the relatively minor average temperature increases from long-term warming (Table 4). Thus, by cooling and humidifying lowland areas, newly formed lowland forests have seemingly enabled the “montane” anoles to follow the forests spreading downslope from montane into lowland habitats (Figure 2). Downslope range movement of some species elsewhere also appear related to habitat modifications (Lenoir et al., 2010).

**Table 4:** Difference in mean annual temperature and precipitation at NOAA weather stations in Puerto Rico, 1952-1977 vs 1991-2015, with *p* values and degrees of freedom for a Welch’s two-sample t-test. *‡* = stations in urban areas.

| Station | Years | Mean<br>Temp<br>°C | Temp<br>shift<br>$\Delta_t$ | $p(\Delta_t = 0)$ | Mean<br>Precip<br>(cm) | Precip<br>shift<br>$\Delta_p$ | $p(\Delta_p = 0)$ |
| --- | --- | --- | --- | --- | --- | --- | --- |
| Borinquen Airport‡ | 1952-1977 | 25.42 | 0.70 | 0.035 | 137.82 | -14.38 | 0.27 |
|  | 1991-2015 | 26.12 |  |  | 123.44 |  |  |
| Corozal Substation | 1952-1977 | 24.31 | 0.59 | 0.003 | 190.09 | -49.32 | 0.01 |
|  | 1991-2015 | 24.90 |  |  | 140.77 |  |  |
| Dos Bocas | 1952-1977 | 25.69 | -0.47 | 0.001 | 194.79 | -20.54 | 0.12 |
|  | 1991-2015 | 25.22 |  |  | 174.26 |  |  |
| Lajas Substation | 1952-1977 | 24.93 | 0.53 | 0.005 | 100.28 | 15.50 | 0.06 |
|  | 1991-2015 | 25.46 |  |  | 115.78 |  |  |
| Manati 2 E | 1952-1977 | 25.33 | 0.12 | 0.240 | 145.76 | 6.73 | 0.64 |
|  | 1991-2015 | 25.45 |  |  | 152.49 |  |  |
| Ponce 4 E | 1952-1977 | 26.09 | 0.34 | 0.056 | 80.60 | 10.18 | 0.31 |
|  | 1991-2015 | 26.44 |  |  | 90.79 |  |  |
| Rio Piedras‡ | 1952-1977 | 25.47 | 0.35 | 0.068 | 164.56 | -50.15 | 0.00 |
|  | 1991-2015 | 25.83 |  |  | 114.41 |  |  |
| Roosevelt Roads‡ | 1952-1977 | 26.68 | 0.31 | 0.110 | 135.47 | -13.51 | 0.41 |
|  | 1991-2015 | 26.99 |  |  | 121.96 |  |  |

But are downward movements of *Anolis* novel invasions, or are they re-invasions in to ancestral zones that were previously forested before agriculture? Paleo-botanical studies find that lowland Puerto Rico was heavily forested prior to 4,800 years BP, when humans first settled in Puerto Rico and began clearing forests and collecting hardwoods for tools, furniture, and ceremonial objects (Rivera-Collazo, 2015). Europeans invaded in the late 15th Century, introducing large-scale agriculture, which accelerated forest clearing, even in remote upland areas (Rivera-Collazo, 2015). Thus, when biologists first began studying *Anolis* in Puerto Rico in early to mid-20th Century, most surviving forests were in montane or karst refugia (Figure 1, Figure 3) (Helmer et al., 2008; Lugo and Helmer, 2004). It is therefore not surprising that biologists initially (if incorrectly) categorized forest-restricted species such as *A. gundlachi* as “montane”. Thus, this is another example of how biogeographic interpretation is sometimes confounded by the “ghosts” of past human activities (Williams, 1972).

We propose that *A. gundlachi* was – prior to European invasions – widespread in both lowland and upland forests, but was later extirpated from lowland (and many upland) areas when forests there were cleared for agriculture. This anole currently occupies lowland forests, even near sea level, despite its low-temperature physiology (Huey and Webster, 1976). In fact, *A. gundlachi* in the forest at Carabalí (*∼* 24 m) does not show elevated corticosterone titers or reduced reproduction in the summer (*Luisa Otero*, *pers. comm.*), as might be expected if these anoles were heat stressed in lowland areas.

*Anolis gundlachi* not only survives in invaded lowland forests, but may even have the capacity to replace *A. cristatellus* there. At Campamento Eliza Colberg and vicinity in 1972 (≈100m), (Gorman and Licht, 1974; Huey, 1974; Huey and Webster, 1976), *A. cristatellus* occurred in both open and forest habitats; and *A. gundlachi* was not observed. Today *A. cristatellus* –but not A. gundlachi – is found the open (N = 74), but *A. gundlachi* is much more common in the forest than is *A. cristatellus* (82 vs. 9 individuals). We suspect similar turnovers have occurred in other new lowland forests, at least those with corridors to upland forest fragments. The ecological mechanism underlying this replacement is unknown, but might reflect a demographic advantage for *A. gundlachi* in winter: 61.45% of female *A. gundlachi* in the forest at Carabalí were gravid in winter (Jan – Feb, N = 68), whereas essentially 0% of *A. cristatellus* in the forest at Pta. Salinas (sea level) were gravid during that season (Otero et al., 2015). Thus, *A. gundlachi* might be better suited to lowland forests than is *A. cristatellus*, at least in cool seasons.

This ecological replacement hypothesis is experimentally testable by introducing (or “reintroducing”) *A. gundlachi* to lowland forest sites (e.g., at Pta. Salinas) that are isolated from other forest patches and where *A. cristatellus* is currently the only trunk-ground anole. Replacement by *A. gundlachi* would support this hypothesis.

Although our primary focus has been on the lowland shifts by montane species, we did find that lowland Anolis (*A. cristatellus*, *A. pulchellus*, *A. stratulus*) though still common at lowland sites are now found at higher elevations than recorded in early collections. Although this shift might reflect a warming-promoted invasion of highland sites, it might indicate a response to newly opened habitats in upland areas following massive forest blow-downs caused by two major hurricanes (Hugo in 1989, Georges in 1998). Such blowdowns lead to warmer and drier operative conditions and could therefore transiently favor lowland species, at least until the forests recover.

## 7 Concluding Remarks

Our study underscores two obvious – but often overlooked – lessons for studies of responses to climate change:

(1) Contemporary biogeographic patterns can reflect real but often well-hidden influences of past human intervention (Nogués-Bravo et al., 2008; Rivera-Collazo, 2015; Williams, 1983). For example, the classic assignment of *A. gundlachi* as a “montane forest” species is almost certainly a historical artifact. When biologists began studying these forest lizards, most remnant forests were restricted to the mountains (Figure 3), as lowland forests had long before been cleared for agriculture (Helmer et al., 2008; Lugo and Helmer, 2004). The low-temperature (and high water loss) physiology of *A. gundlachi* in particular reinforced the assumption that this species should be restricted to cool upland forests (Gorman and Hillman, 1977; Heatwole et al., 1969; Huey and Webster, 1976). However, our discovery of *A. gundlachi* in newly regenerated lowland forests suggests that this species was likely once native to lowland – as well as upland – forests prior to human disturbance. Thus *A. gundlachi* is more properly considered a “forest” species. The hypothesis that this species was native to lowland forests is potentially testable by fossil evidence; and the hypothesis that lowland populations are recent invaders is potentially testable by genetic analysis (Peter and Slatkin, 2013).

(2) Other studies have also emphasized that impacts of climate change must recognize that climate may not be the most influential environmental factor driving species range shifts (Lenoir et al., 2010; Lenoir and Svenning, 2015; Nogués-Bravo et al., 2008; Seabra et al., 2015; Battey, 2019; Guo et al., 2018; Santos et al., 2017). In the present case, the magnitude of local cooling caused by a regenerated lowland forest swamps the minor (average) temperature increases caused so far by anthropogenic warming. Moreover, regenerating forests at low elevations dramatically expands the area suitable for “montane” lizards (though this pattern may eventually reverse given sufficient warming; Barros et al. (2014); Nowakowski et al. (2017)). Perhaps more importantly, temperature and humidity are just two axes of variation in the complex ecosystem in which montane *Anolis* are embedded. Shifts in food availability, disease, and predation (among many other factors) occur along with the thermal and hydric impacts of lowland forest regeneration (Buckley, 2013; Otero et al., 2019), and these factors all affect the expansion of *Anolis* populations at low elevation.

The reforestation of Puerto Rico should not be interpreted as a general model for the future of tropical forests, as human-driven deforestation is clearly the norm in most tropical regions. Moreover, forest coverage and age in Puerto Rico are also strongly affected by frequent hurricanes, which dynamically if transiently alter the ecology and distributions of plants and animals (Lugo et al., 1981; Uriarte et al., 2009). In any case, when tropical climate warming (Battisti and Naylor, 2009) combines with deforestation – caused either by natural (e.g., from hurricanes) or anthropogenic forces – local operative temperatures will increase substantially and may harm all but the most heat tolerant species (Colwell et al., 2008; Frishkoff et al., 2015; Sunday et al., 2010).

## 8 Acknowledgements

We thank Héctor Álvarex Pérez for help in the conceptual development of this project. Research was supported by collaborative grants from NSF to RBH (1038016), PEH (1038012), BL (1038013), and HJA and PAB (1038014). We thank M. A. García, A. Gunderson, M. Leal, A. E. Lugo, L. Revell, J. Rodríguez-Robles, and R. Thomas for help and discussion, and A. Diaz for permission to work at Carabalí. Many thanks to Andrew Kern for advice, support for publication costs, and for creating a supportive lab environment that let CJB get this thing in shape for submission. Research was conducted under permits from the Departamento de Recursos Naturales y Ambientales (Puerto Rico) and with approved Institutional Animal Care and Use Committee protocols from the University of Washington and the University of Puerto Rico.

## 10 Authors’ Contributions

All authors contributed to conceptual development of the initial elevation study, and RH and CJB conceived the retrospective study design for specimen data. CJB and RH drafted the initial text with input from LO, GG, PH, BL, AG, and PAB. CJB conducted statistical analyses, georeferenced localities, and prepared figures. RH, LO, GG, PH, BL, AG, PB, and HP contributed field data, suggested many improvements to the manuscript during drafting, and assisted with interpretation of results.

## 11 Supplementary Figures and Tables

**Table S1:** MAP parameter estimates for the linear model, by species.

|  |  | Mean | StdDev | 2.50% | 97.50% |
| --- | --- | --- | --- | --- | --- |
| *evermanni* | $A_i$ | 398.34 | 18.08 | 362.91 | 433.77 |
| | $E_i$ | -17.04 | 5.54 | -27.89 | -6.19 |
| | $D_i$ | -66.32 | 22.7 | -114.82 | -19.01 |
| | $\sigma_i$ | 0.91 | 0.02 | 0.86 | 0.96 |
| | $\log(\text{Likelihood})$ | -4863.82 | | | |
| *gundlachi* | $A_i$ | 489.78 | 6.07 | 477.89 | 501.68 |
| | $E_i$ | 22.76 | 6.18 | 10.65 | 34.86 |
| | $D_i$ | -171.08 | 10.68 | -193.41 | -149.56 |
| | $\sigma_i$ | 0.43 | 0.01 | 0.42 | 0.45 |
| | $\log(\text{Likelihood})$ | -11509.46 | | | |
| *krugi* | $A_i$ | 354.37 | 8.05 | 338.6 | 370.15 |
| | $E_i$ | 3.19 | 5.64 | -7.87 | 14.24 |
| | $D_i$ | -146.33 | 12.14 | -169.96 | -122.36 |
| | $\sigma_i$ | 0.74 | 0.01 | 0.71 | 0.76 |
| | $\log(\text{Likelihood})$ | -10070.34 | | | |
| *cristatellus* | $A_i$ | 29.84 | 1.37 | 27.15 | 32.52 |
| | $E_i$ | -33.51 | 7.13 | -47.49 | -19.53 |
| | $D_i$ | 73.44 | 8.15 | 57.31 | 88.96 |
| | $\sigma_i$ | 1.77 | 0.03 | 1.72 | 1.82 |
| | $\log(\text{Likelihood})$ | -13162.95 | | | |
| *pulchellus* | $A_i$ | 32.47 | 1.82 | 28.9 | 36.04 |
| | $E_i$ | -20.35 | 6.13 | -32.37 | -8.33 |
| | $D_i$ | 26.11 | 7.01 | 12.37 | 39.85 |
| | $\sigma_i$ | 1.89 | 0.03 | 1.83 | 1.96 |
| | $\log(\text{Likelihood})$ | -9751.37 | | | |
| *stratulus* | $A_i$ | 57.92 | 3.72 | 50.62 | 65.22 |
| | $E_i$ | -19.66 | 5.73 | -30.9 | -8.42 |
| | $D_i$ | 17.55 | 8.6 | 0.92 | 33.87 |
| | $\sigma_i$ | 1.55 | 0.04 | 1.48 | 1.62 |
| | $\log(\text{Likelihood})$ | -5371.34 | | | |

**Figure S1:**
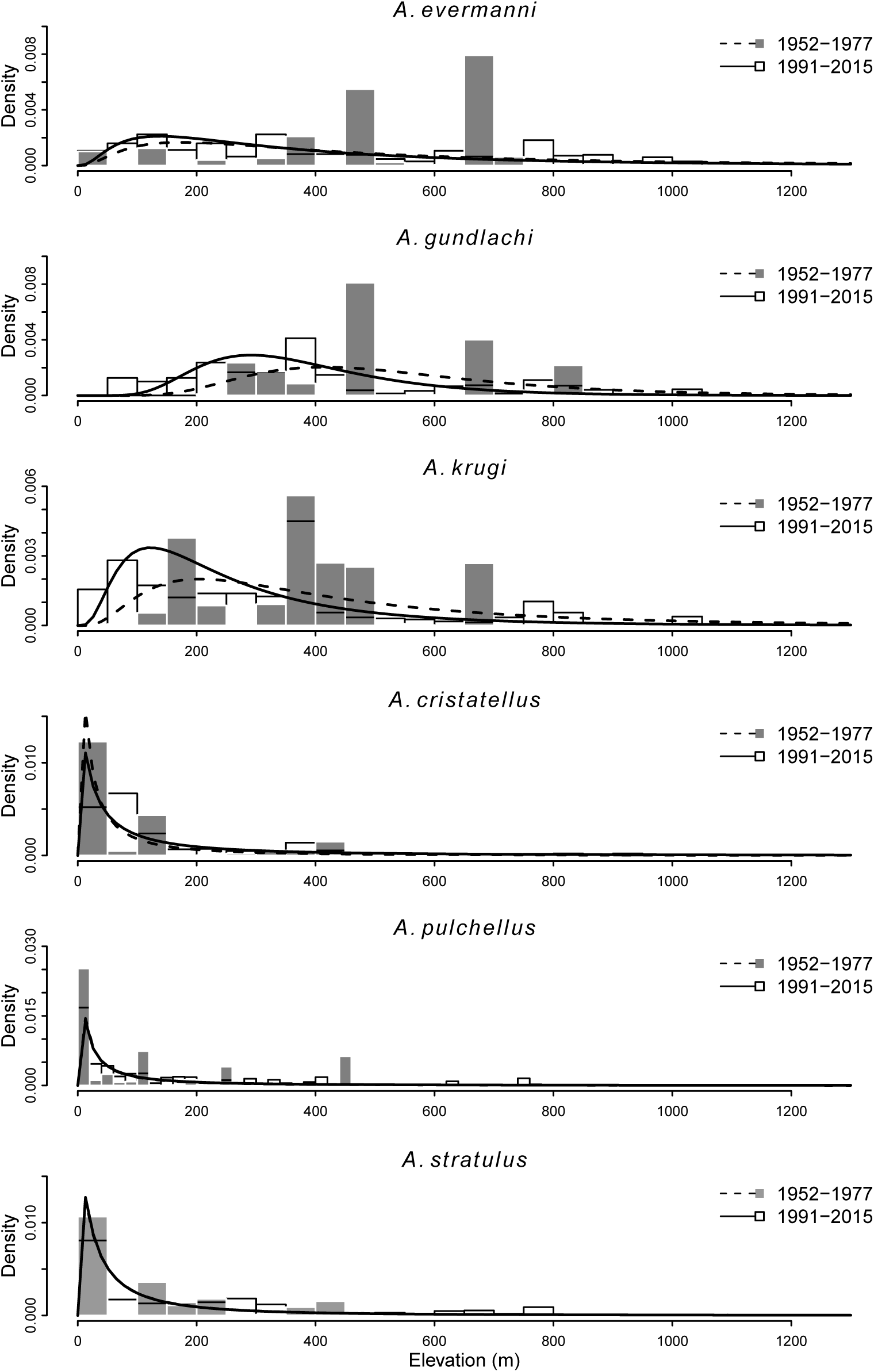
Linear model fits. Histograms are empirical density of species reports by eleva-tion and curves are the corresponding log-normal distributions fit by the MAP analysis.

**Figure S2:**
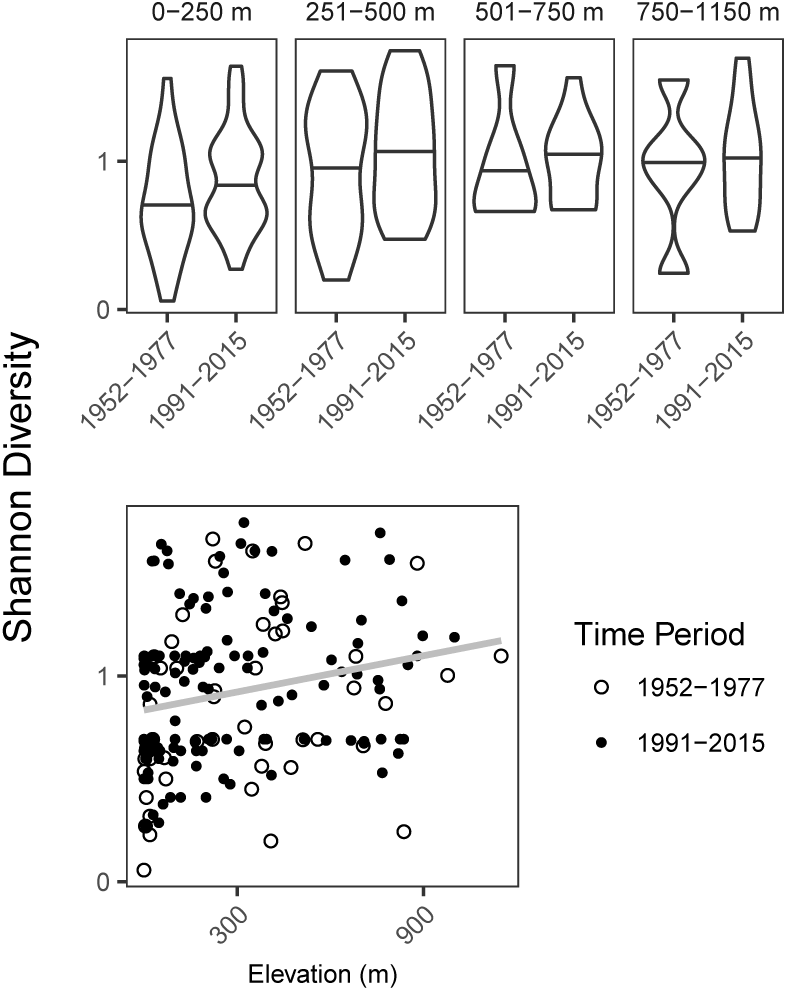
(A) Pairwise differences in Shannon diversity per locality cluster across time periods in elevation bins. No pairwise differences are statistically significant. (B) linear regression of diversity as a function of locality elevation. ANOVA analyses found that incorporating time period did not significantly improve model fit so a single regression across all locations and time periods is shown.

**Figure S3:**
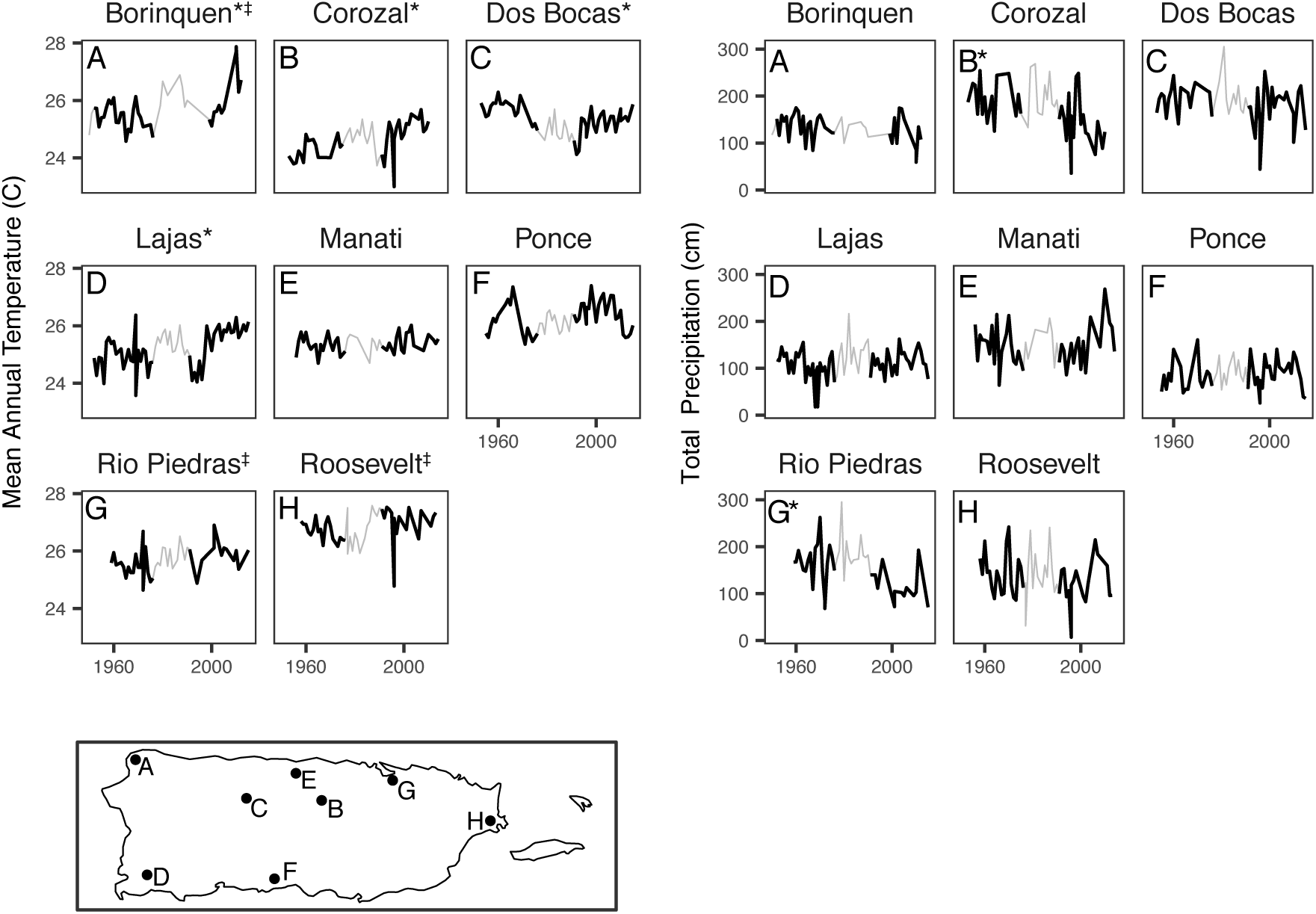
Mean annual temperatures (left) and total precipitation (right) for weather stations with at least 40 years of complete records on Puerto Rico. * stations with significant difference in means across time periods (dark lines), *‡* station is in an urban area. Bottom: station locations.

**Figure S4:**
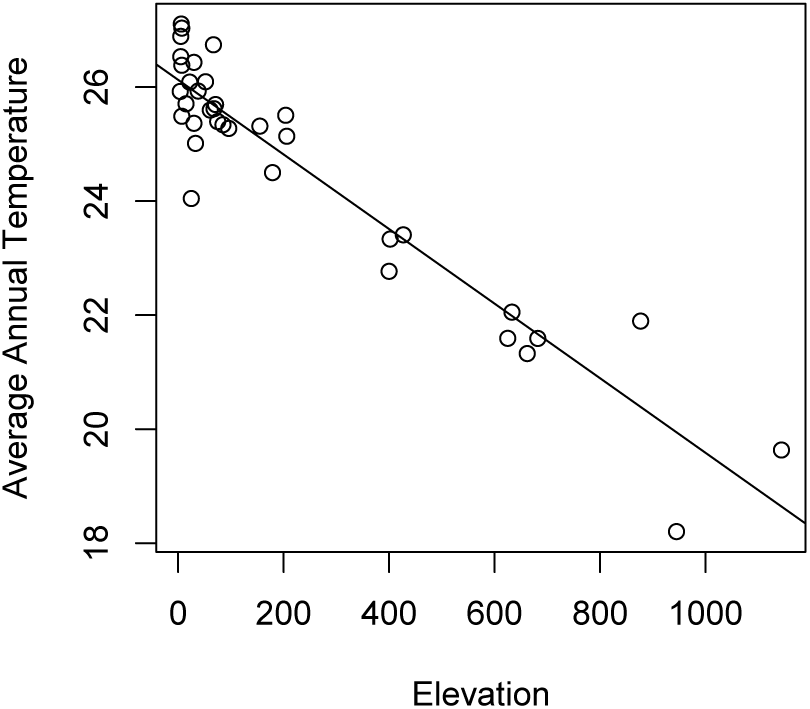
Relationship between temperature and altitude for 16 weather stations on Puerto Rico (4-1144 meters elevation). The slope of the linear model is −6.541°C/km and R2 is 0.887.

## References

Aide, T. M., Zimmerman, J. K., Rosario, M. and Marcano, H. (1996), ‘Forest recovery in abandoned cattle pastures along an elevational gradient in northeastern puerto rico’, Biotropica pp. 537–548.

Álvarez-Berríos, N., Redo, D., Aide, T., Clark, M. and Grau, R. (2013), ‘Land change in the greater antilles between 2001 and 2010’, Land 2(2), 81–107.

Angert, A. L., LaDeau, S. L. and Ostfeld, R. S. (2013), ‘Climate change and species interactions: ways forward’, Annals of the New York Academy of Sciences 1297(1), 1–7.

Bakken, G. S. (1992), ‘Measurement and application of operative and standard operative temperatures in ecology’, American Zoologist 32(2), 194–216.

Barros, V., Field, C., Dokke, D., Mastrandrea, M., Mach, K., Bilir, T., Chatterjee, M., Ebi, K., Estrada, Y., Genova, R. et al. (2014), ‘Climate change |v2014: |pimpacts, adaptation, and vulnerability-part b: regional aspects-contribution of working group ii to the fifth assessment report of the intergovernmental panel on climate change’.

Bastable, H., Shuttleworth, W. J., Dallarosa, R., Fisch, G. and Nobre, C. A. (1993), ‘Observations of climate, albedo, and surface radiation over cleared and undisturbed amazonian forest’, International Journal of Climatology 13(7), 783–796.

Battey, C. J. (2019), ‘Ecological release of the anna’s hummingbird during a northern range expansion’, The American Naturalist 194(3), 306–315.

URL: https://doi.org/10.1086/704249

Battisti, D. S. and Naylor, R. L. (2009), ‘Historical warnings of future food insecurity with unprecedented seasonal heat’, Science 323(5911), 240–244.

Beck, J. and Schwanghart, W. (2010), ‘Comparing measures of species diversity from incomplete inventories: an update’, Methods in Ecology and Evolution 1(1), 38–44.

Bivand, R. and Rundel, C. (2013), ‘rgeos: interface to geometry engine-open source (geos)’, R package version 0.3-2.

Buckley, L. B. (2013), ‘Get real: putting models of climate change and species interactions in practice’, Annals of the New York Academy of Sciences 1297(1), 126–138.

Burrowes, P. A., Joglar, R. L. and Green, D. E. (2004), ‘Potential causes for amphibian declines in puerto rico’, Herpetologica 60(2), 141–154.

Camacho, A., Van den Brooks, J. M., Riley, A., Telemeco, R. S. and Angilletta Jr, M. J. (2018), ‘Oxygen supply did not affect how lizards responded to thermal stress’, Integrative Zoology 13(4), 428–436.

Canham, C. D., Thompson, J., Zimmerman, J. K. and Uriarte, M. (2010), ‘Variation in susceptibility to hurricane damage as a function of storm intensity in puerto rican tree species’, Biotropica 42(1), 87–94.

Chen, I.-C., Hill, J. K., Ohlemüller, R., Roy, D. B. and Thomas, C. D. (2011), ‘Rapid range shifts of species associated with high levels of climate warming’, Science 333(6045), 1024–1026.

Colwell, R. K., Brehm, G., Cardel ú s, C. L., Gilman, A. C. and Longino, J. T. (2008), ‘Global warming, elevational range shifts, and lowland biotic attrition in the wet tropics’, science 322(5899), 258–261.

Comarazamy, D. E. and Gonz á lez, J. E. (2011), ‘Regional long-term climate change (1950–2000) in the midtropical atlantic and its impacts on the hydrological cycle of puerto rico’, Journal of Geophysical Research: Atmospheres 116(D21).

Davis, A. J., Jenkinson, L. S., Lawton, J. H., Shorrocks, B. and Wood, S. (1998), ‘Making mistakes when predicting shifts in species range in response to global warming’, Nature 391(6669), 783.

Deutsch, C. A., Tewksbury, J. J., Huey, R. B., Sheldon, K. S., Ghalambor, C. K., Haak, D. C. and Martin, P. R. (2008), ‘Impacts of climate warming on terrestrial ectotherms across latitude’, Proceedings of the National Academy of Sciences 105(18), 6668–6672.

Freeman, B. G. and Freeman, A. M. C. (2014), ‘Rapid upslope shifts in new guinean birds illustrate strong distributional responses of tropical montane species to global warming’, Proceedings of the National Academy of Sciences 111(12), 4490–4494.

Frishkoff, L. O., Hadly, E. A. and Daily, G. C. (2015), ‘Thermal niche predicts tolerance to habitat conversion in tropical amphibians and reptiles’, Global change biology 21(11), 3901–3916.

Geiger, R., Aron, R. H. and Todhunter, P. (2009), The climate near the ground, Rowman & Little-field.

Gorman, G. C. and Hillman, S. (1977), ‘Physiological basis for climatic niche partitioning in two species of puerto rican *Anolis* (reptilia, lacertilia, iguanidae)’, Journal of Herpetology pp. 337–340.

Gorman, G. C. and Licht, P. (1974), ‘Seasonality in ovarian cycles among tropical *Anolis* lizards’, Ecology 55(2), 360–369.

Grau, H. R., Aide, T. M., Zimmerman, J. K., Thomlinson, J. R., Helmer, E. and Zou, X. (2003), ‘The ecological consequences of socioeconomic and land-use changes in postagriculture puerto rico’, BioScience 53(12), 1159–1168.

Gunderson, A. R. and Leal, M. (2012), ‘Geographic variation in vulnerability to climate warming in a tropical caribbean lizard’, Functional Ecology 26(4), 783–793.

Gunderson, A. R., Mahler, D. L. and Leal, M. (2016), ‘A functional analysis of the contribution of climatic niche divergence to adaptive radiation’, bioRxiv p. 069294.

Guo, F., Lenoir, J. and Bonebrake, T. C. (2018), ‘Land-use change interacts with climate to determine elevational species redistribution’, Nature communications 9(1), 1315.

Harmsen, E. W., Miller, N. L., Schlegel, N. J. and Gonzalez, J. E. (2009), ‘Seasonal climate change impacts on evapotranspiration, precipitation deficit and crop yield in puerto rico’, Agricultural water management 96(7), 1085–1095.

Heatwole, H. (1970), ‘Thermal ecology of the desert dragon amphibolurus inermis’, Ecological Monographs 40(4), 425–457.

Heatwole, H., Lin, T.-H., Villal ón, E., Muñiz, A. and Matta, A. (1969), ‘Some aspects of the thermal ecology of puerto rican anoline lizards’, Journal of Herpetology pp. 65–77.

Helmer, E., Brandeis, T. J., Lugo, A. E. and Kennaway, T. (2008), ‘Factors influencing spatial pattern in tropical forest clearance and stand age: Implications for carbon storage and species diversity’, Journal of Geophysical Research: Biogeosciences 113(G2).

Helmus, M. R., Mahler, D. L. and Losos, J. B. (2014), ‘Island biogeography of the anthropocene’, Nature 513(7519), 543.

Hertz, P., Arce-Hernandez, A., Ramirez-Vazquez, J., Tirado-Rivera, W. and Vazquez-Vives, L. (1979), ‘Geographical variation of heat sensitivity and water loss rates in the tropical lizard, Anolis gundlachi’, Comparative Biochemistry and Physiology Part A: Physiology 62(4), 947–953.

Hertz, P. E. (1981), ‘Adaptation to altitude in two west indian anoles (reptilia’, Journal of Zoology 195(1), 25–37.

Hertz, P. E. (1992), ‘Temperature regulation in puerto rican *Anolis* lizards: a field test using null hypotheses’, Ecology 73(4), 1405–1417.

Hertz, P. E., Arima, Y., Harrison, A., Huey, R. B., Losos, J. B. and Glor, R. E. (2013), ‘Asynchronous evolution of physiology and morphology in *Anolis* lizards’, Evolution 67(7), 2101–2113.

Hijmans, R. J. (2015), ‘Geographic data analysis and modeling [r package raster version 2.8-19]’.

Holm, S. (1979), ‘A simple sequentially rejective multiple test procedure’, Scandinavian journal of statistics pp. 65–70.

Huey, R. B. (1974), ‘Behavioral thermoregulation in lizards: importance of associated costs’, Science 184(4140), 1001–1003.

Huey, R. B. (1978), ‘Latitudinal pattern of between-altitude faunal similarity: mountains might be “higher” in the tropics’, The American Naturalist 112(983), 225–229.

Huey, R. B., Deutsch, C. A., Tewksbury, J. J., Vitt, L. J., Hertz, P. E., Álvarez Pérez, H. J. and Garland Jr, T. (2009), ‘Why tropical forest lizards are vulnerable to climate warming’, Proceedings of the Royal Society B: Biological Sciences 276(1664), 1939–1948.

Huey, R. B. and Webster, T. P. (1976), ‘Thermal biology of *Anolis* lizards in a complex fauna: the christatellus group on puerto rico’, Ecology 57(5), 985–994.

Janzen, D. H. (1967), ‘Why mountain passes are higher in the tropics’, The American Naturalist 101(919), 233–249.

Jennings, L. N., Douglas, J., Treasure, E. and González, G. (2014), ‘Climate change effects in el yunque national forest, puerto rico, and the caribbean region’, Gen. Tech. Rep. SRS-GTR-193. Asheville, NC: USDA-Forest Service, Southern Research Station. 47 p. 193, 1–47.

Karl, T. R., Arguez, A., Huang, B., Lawrimore, J. H., McMahon, J. R., Menne, M. J., Peterson, T. C., Vose, R. S. and Zhang, H.-M. (2015), ‘Possible artifacts of data biases in the recent global surface warming hiatus’, Science 348(6242), 1469–1472.

Kaspari, M., Clay, N. A., Lucas, J., Yanoviak, S. P. and Kay, A. (2015), ‘Thermal adaptation generates a diversity of thermal limits in a rainforest ant community’, Global Change Biology 21(3), 1092–1102.

Leal, M. and Fleishman, L. J. (2002), ‘Evidence for habitat partitioning based on adaptation to environmental light in a pair of sympatric lizard species’, Proceedings of the Royal Society of London. Series B: Biological Sciences 269(1489), 351–359.

Lenoir, J., Gégout, J.-C., Guisan, A., Vittoz, P., Wohlgemuth, T., Zimmermann, N. E., Dullinger, S., Pauli, H., Willner, W. and Svenning, J.-C. (2010), ‘Going against the flow: potential mechanisms for unexpected downslope range shifts in a warming climate’, Ecography 33(2), 295–303.

Lenoir, J. and Svenning, J.-C. (2015), ‘Climate-related range shifts–a global multidimensional synthesis and new research directions’, Ecography 38(1), 15–28.

Linck, E., Bridge, E. S., Duckles, J. M., Navarro-Sigüenza, A. G. and Rohwer, S. (2016), ‘Assessing migration patterns in *Passerina ciris* using the world’s bird collections as an aggregated resource’, PeerJ 4, e1871.

Lister, B. C. (1981), ‘Seasonal niche relationships of rain forest anoles’, Ecology 62(6), 1548–1560.

Lister, B. C. and Garc ía, A. (2018), ‘Climate-driven declines in arthropod abundance restructure a rainforest food web’, Proceedings of the National Academy of Sciences 115(44), E10397–E10406.

Losos, J. B. (2011), Lizards in an evolutionary tree: ecology and adaptive radiation of anoles, Vol. 10, Univ of California Press.

Lugo, A. E. and Helmer, E. (2004), ‘Emerging forests on abandoned land: Puerto rico’s new forests’, Forest Ecology and Management 190(2-3), 145–161.

Lugo, A. E., Schmidt, R. and Brown, S. (1981), ‘Tropical forests in the caribbean.’, Ambio. Stockholm 10(6), 318–324.

McCain, C. M. and Colwell, R. K. (2011), ‘Assessing the threat to montane biodiversity from discordant shifts in temperature and precipitation in a changing climate’, Ecology letters 14(12), 1236–1245.

McElreath, R. (2012), ‘rethinking: Statistical rethinking book package’, R package version 1.

Moritz, C., Patton, J. L., Conroy, C. J., Parra, J. L., White, G. C. and Beissinger, S. R. (2008), ‘Impact of a century of climate change on small-mammal communities in yosemite national park, usa’, Science 322(5899), 261–264.

Morueta-Holme, N., Engemann, K., Sandoval-Acuña, P., Jonas, J. D., Segnitz, R. M. and Svenning, J.-C. (2015), ‘Strong upslope shifts in chimborazo’s vegetation over two centuries since humboldt’, Proceedings of the National Academy of Sciences 112(41), 12741–12745.

Murphy, D. J., Hall, M. H., Hall, C. A., Heisler, G. M., Stehman, S. V. and Anselmi-Molina, C. (2011), ‘The relationship between land cover and the urban heat island in northeastern puerto rico’, International Journal of Climatology 31(8), 1222–1239.

Méndez-Lázaro, P., Mart ínez-Sánchez, O., Méndez-Tejeda, R., Rodr íguez, E., Morales, E. and Schmitt-Cortijo, N. (2015), ‘Extreme heat events in san juan puerto rico: Trends and variability of unusual hot weather and its possible effects on ecology and society’, Journal of Climatology and Weather Forecasting 3(2), 1–7.

Méndez-Tejeda, R. (2017), ‘Increase in the number of hot days for decades in puerto rico 1950-2014’, Environment and Natural Resources Research 7(3), 16–26.

Nogu é s-Bravo, D., Ara ú jo, M., Romdal, T. and Rahbek, C. (2008), ‘Scale effects and human impact on the elevational species richness gradients’, Nature 453(7192), 216.

Nowakowski, A. J., Watling, J. I., Whitfield, S. M., Todd, B. D., Kurz, D. J. and Donnelly, M. A. (2017), ‘Tropical amphibians in shifting thermal landscapes under land-use and climate change’, Conservation biology 31(1), 96–105.

Oke, T. R. (1982), ‘The energetic basis of the urban heat island’, Quarterly Journal of the Royal Meteorological Society 108(455), 1–24.

Otero, L. M., Huey, R. B. and Gorman, G. C. (2015), ‘A few meters matter: local habitats drive reproductive cycles in a tropical lizard’, The American Naturalist 186(3), E72–E80.

Otero, L., Schall, J. J., Cruz, V., Aaltonen, K. and Acevedo, M. A. (2019), ‘The drivers and consequences of unstable plasmodium dynamics: a long-term study of three malaria parasite species infecting a tropical lizard’, Parasitology 146(4), 453–461.

Otero López, L. M. and Sabat, A. M. (2015), ‘Reproductive phenology, fecundity, survival and growth of puerto rican anolis lizards in the context of climate warming’.

Parmesan, C. (2006), ‘Ecological and evolutionary responses to recent climate change’, Annu. Rev. Ecol. Evol. Syst. 37, 637–669.

Patz, J. A., Martens, W., Focks, D. A. and Jetten, T. H. (1998), ‘Dengue fever epidemic potential as projected by general circulation models of global climate change.’, Environmental health perspectives 106(3), 147–153.

Peter, B. M. and Slatkin, M. (2013), ‘Detecting range expansions from genetic data’, Evolution 67(11), 3274–3289.

URL: https://onlinelibrary.wiley.com/doi/abs/10.1111/evo.12202

Polato, N. R., Gill, B. A., Shah, A. A., Gray, M. M., Casner, K. L., Barthelet, A., Messer, P. W., Simmons, M. P., Guayasamin, J. M., Encalada, A. C. et al. (2018), ‘Narrow thermal tolerance and low dispersal drive higher speciation in tropical mountains’, Proceedings of the National Academy of Sciences 115(49), 12471–12476.

Pounds, J. A., Fogden, M. P. and Campbell, J. H. (1999), ‘Biological response to climate change on a tropical mountain’, Nature 398(6728), 611.

Rand, A. S. (1964), ‘Ecological distribution in anoline lizards of puerto rico’, Ecology 45(4), 745–752.

Raxworthy, C. J., Pearson, R. G., Rabibisoa, N., Rakotondrazafy, A. M., Ramanamanjato, J.-B., Raselimanana, A. P., Wu, S., Nussbaum, R. A. and Stone, D. A. (2008), ‘Extinction vulnerability of tropical montane endemism from warming and upslope displacement: a preliminary appraisal for the highest massif in madagascar’, Global Change Biology 14(8), 1703–1720.

Rivera-Collazo, I. C. (2015), ‘Por el camino verde: Long-term tropical socioecosystem dynamics and the anthropocene as seen from puerto rico’, The Holocene 25(10), 1604–1611.

Rivero, J. A. (1998), Amphibians and reptiles of Puerto Rico, La Editorial, UPR.

Rodr íguez-Robles, J. A., Leal, M. and Losos, J. B. (2005), ‘Habitat selection by the puerto rican yellow-chinned anole, Anolis gundlachi’, Canadian journal of zoology 83(7), 983–988.

Rohwer, S., Rohwer, V. G., Peterson, A. T., Navarro-Sigüenza, A. G. and English, P. (2012), ‘Assessing migratory double breeding through complementary specimen densities and breeding records’, The Condor 114(1), 1–14.

Santos, M. J., Smith, A. B., Thorne, J. H. and Moritz, C. (2017), ‘The relative influence of change in habitat and climate on elevation range limits in small mammals in yosemite national park, california, usa’, Climate Change Responses 4(1), 7.

Schmidt, K. P. (1918), ‘Contributions to the herpetology of porto rico’, Annals of the New York Academy of Sciences 28(1), 167–200.

Schoener, T. W. (1971), Structural habitats of West Indian Anolis lizards, Harvard University, Museum of Comparative Zoology.

Schoener, T. W. and Schoener, A. (1971), ‘Structural habitats of west indian *Anolis* lizards ii. puerto rican uplands’, Breviora 35, 1–30.

Schwartz, A. and Henderson, R. W. (1991), Amphibians and reptiles of the West Indies: descriptions, distributions, and natural history, University Press of Florida.

Seabra, R., Wethey, D. S., Santos, A. M. and Lima, F. P. (2015), ‘Understanding complex biogeographic responses to climate change’, Scientific reports 5, 12930.

Shannon, C. E. (1948), ‘A mathematical theory of communication’, Bell system technical journal 27(3), 379–423.

Sunday, J. M., Bates, A. E. and Dulvy, N. K. (2010), ‘Global analysis of thermal tolerance and latitude in ectotherms’, Proceedings of the Royal Society B: Biological Sciences 278(1713), 1823–1830.

Taubert, F., Fischer, R., Groeneveld, J., Lehmann, S., Müller, M. S., Rödig, E., Wiegand, T. and Huth, A. (2018), ‘Global patterns of tropical forest fragmentation’, Nature 554(7693), 519.

Team, R. C. (2014), ‘R: A language and environment for statistical computing [internet]. 2013’, Disponible sur: http://www.R-project.org.

Tingley, M. W. and Beissinger, S. R. (2009), ‘Detecting range shifts from historical species occurrences: new perspectives on old data’, Trends in ecology & evolution 24(11), 625–633.

Uriarte, M., Canham, C. D., Thompson, J., Zimmerman, J. K., Murphy, L., Sabat, A. M., Fetcher, N. and Haines, B. L. (2009), ‘Natural disturbance and human land use as determinants of tropical forest dynamics: results from a forest simulator’, Ecological Monographs 79(3), 423–443.

USGS (2016), ‘The national map 3d elevation program’.

URL: http://nationalmap.gov/3DEP/3depprodserv.html

Waide, R. B., Comarazamy, D. E., Gonz á lez, J. E., Hall, C. A., Lugo, A. E., Luvall, J. C., Murphy, D. J., Ortiz-Zayas, J. R., Ram írez-Beltran, N. D., Scatena, F. N. et al. (2013), ‘Climate variability at multiple spatial and temporal scales in the luquillo mountains, puerto rico’.

URL: https://data.fs.usda.gov/research/pubs/iitf/BC_EB54-12_Waide_et_al.pdf

Wake, D. and Lynch, J. (1976), ‘The distribution, ecology, and evolutionary history of plethodontid salamanders in tropical america.’, Natural History 25, 1–65.

Walker, L. R. (1991), ‘Tree damage and recovery from hurricane hugo in luquillo experimental forest, puerto rico’, Biotropica pp. 379–385.

Wieczorek, J. (2001), ‘Manis: Georeferencing guidelines’, University of California, Berkeley– MaNIS.[cited 26 January 2004] 37.

Williams, E. E. (1972), The origin of faunas. evolution of lizard congeners in a complex island fauna: a trial analysis, in ‘Evolutionary biology’, Springer, pp. 47–89.

Williams, E. E. (1983), ‘Ecomorphs, faunas, island size, and diverse end points in island radiations of anolis’, Lizard ecology: studies of a model organism pp. 327–370.

Yackulic, C. B., Fagan, M. E., Jain, M., Jina, A., Lim, Y., Marlier, M. E., Muscarella, R., Adame, P., DeFries, R. S. and Uriarte, M. (2011), ‘Biophysical and socioeconomic factors associated with forest transitions at multiple spatial and temporal scales’.

